# The RNA helicase DDX5 promotes alveolar rhabdomyosarcoma growth and survival

**DOI:** 10.1101/2020.07.08.194092

**Authors:** Alberto Gualtieri, Valerio Licursi, Chiara Mozzetta

## Abstract

Rhabdomyosarcoma (RMS) is the most common soft-tissue sarcoma of childhood characterized by the inability to exit the proliferative myoblast-like stage. The alveolar fusion positive subtype (FP-ARMS) is the most aggressive and is mainly caused by the expression of PAX3/7-FOXO1 oncoproteins, which are challenging pharmacological targets. Thus, other therapeutic vulnerabilities resulting from gene expression changes are progressively being recognized. Here, we identified the DEAD box RNA helicase 5 (DDX5) as a potential therapeutic target to inhibit FP-ARMS growth. We show that DDX5 is overexpressed in alveolar RMS cells, demonstrating that its depletion drastically decreases FP-ARMS viability and slows tumor growth in xenograft models. Mechanistically, we provide evidence that DDX5 functions upstream the G9a/AKT survival signalling pathway, by modulating G9a protein stability. Finally, we show that G9a interacts with PAX3-FOXO1 and regulates its activity, thus sustaining FP-ARMS myoblastic state. Together, our findings identify a novel survival-promoting loop in FP-ARMS and highlight DDX5 as potential therapeutic target to arrest rhabdomyosarcoma growth.

## Introduction

Rhabdomyosarcomas (RMS) are aggressive childhood cancers representing the most common soft-tissue sarcomas in pediatric population. Around 60% of all children and adolescents diagnosed with RMS are cured by currently available multimodal therapies, including surgery, radiation and conventional chemotherapeutic drugs. However, clinical outcomes for patients with high-risk RMS are still poor, emphasizing the urgency to explore more in depth its molecular underpinnings and to devise new effective therapeutic interventions (1).

Pediatric RMS are typically divided into two major categories: alveolar (ARMS) and embryonal (ERMS). These two types of RMS are clinically and molecularly different. ERMS are more common, histologically resemble embryonic skeletal muscle, arise early in childhood from head, neck and retroperitoneum, and are typically associated with better prognosis. The genetic profile of ERMS is heterogeneous and is associated with activation of various tumor-promoting signaling pathways and/or loss of tumor surveillance. ARMS are most common in older children, predominantly involve trunk and extremities and are generally more aggressive. They typically associate with pathognomonic chromosomal translocations, such as t(2;13) or t(1;13) that result in fusion proteins combining the DNA binding domain of PAX3 or PAX7 with the transcriptional activation domain of FOXO1A, which account for the 60% or 20% ARMS cases, respectively. The remaining 20% of ARMS lack molecular evidence of these translocations and are referred to as fusion-negative ARMS (2).

Alveolar fusion-positive (FP-ARMS) is the most aggressive subtype, associated with frequent metastasis at the time of diagnosis and limited response to treatment, resulting in poor survival rates. FP-RMS cells are addicted to the oncogenic capacity of PAX3/7-FOXO1, which have become very important prognostic markers in the clinics. However, direct targeting of the fusion proteins is still a challenge (3) and other therapeutic vulnerabilities resulting from gene expression changes are being extensively investigated in the last years (4-7). In this context, DEAD box RNA helicases appear appealing candidates as potential therapeutic targets, having been implicated in almost every aspect of RNA metabolism, including transcription, pre-mRNA splicing, ribosome biogenesis, transport, translation, and RNA decay (8).

In normal myogenesis, the DEAD box helicase 5 (DDX5, also known as p68) is needed for proper differentiation, being part of a multitasking complex that together with the steroid nuclear receptor activator (*SRA*) long-non coding RNA, BRG1 and MYOD, promotes transcriptional expression of MYOD-target genes (9). Moreover, DDX5 cooperates with heterogeneous nuclear ribonucleoprotein (hnRNP) to establish specific splicing subprograms in myoblasts along myogenesis (10), highlighting its multimodal actions in shaping the gene expression programs during cell differentiation. Many studies have detected the overexpression of DDX5 in different human malignancies and confirmed its involvement in tumorigenesis, invasion, proliferation and metastasis (11-13). Thus, DDX5 is a potentially valuable diagnostic and prognostic marker in cancer. However, whether DDX5 plays a role in rhabdomyosarcoma pathogenesis has been not addressed yet. Here, we demonstrate that DDX5 is overexpressed in FP-ARMS and that it promotes their survival and growth, both *in vitro* and *in vivo*. Mechanistically, we found that DDX5 interacts and cooperates with the lysine methyltransferase G9a to stabilize PAX3-FOXO1 thus sustaining the myoblastic stage of FP-ARMS.

## Results and Discussion

### DDX5 is overexpressed in rhabdomyosarcoma and sustains FP-ARMS survival

To gain insights into a possible role of DDX5 in RMS, we looked at its expression and epigenetic status through the Integrated Rhabdomyosarcoma database of the St. Jude Children’s Research Hospital (https://pecan.stjude.cloud/proteinpaint/study/RHB2018) (14). Among the 18 chromatin hidden Markov modeling (chromHMM) states (15) identified in the study (14) (**Supplemental Figure. 1A**), DDX5 showed a strong active transcription start site (TSS) (red bars) and an actively transcribed gene body (green bars) in either normal myoblasts and myotubes, and primary ERMS and ARMS samples (**Supplemental Figure. 1B**). Accordingly, *DDX5* RNA levels (*DDX5* FPKM) did not significantly differ among normal and RMS samples (**Figure 1A**, *blue bars*). By contrast, proteomic data revealed that DDX5 protein levels were higher than normal cells in all tested RMS samples (**Figure 1A**, *red bars*), which were also accompanied by an hyperphosphorylated status of DDX5 as compared to normal myoblasts (**Figure 1A**, *circles*). Experimental analysis of DDX5 expression, by quantitative real-time PCR (qRT-PCR) (**Figure 1B)** and Western blot (WB) (**Figure 1C**), on two different FP-ARMS cell lines, RH30 and RH41, confirmed DDX5 overexpression in FP-ARMS, as compared to normal <u>h</u>uman <u>s</u>keletal <u>m</u>uscle <u>m</u>yoblasts (HSMMs), prompting our interest in investigating its functional role.

**Figure 1.**
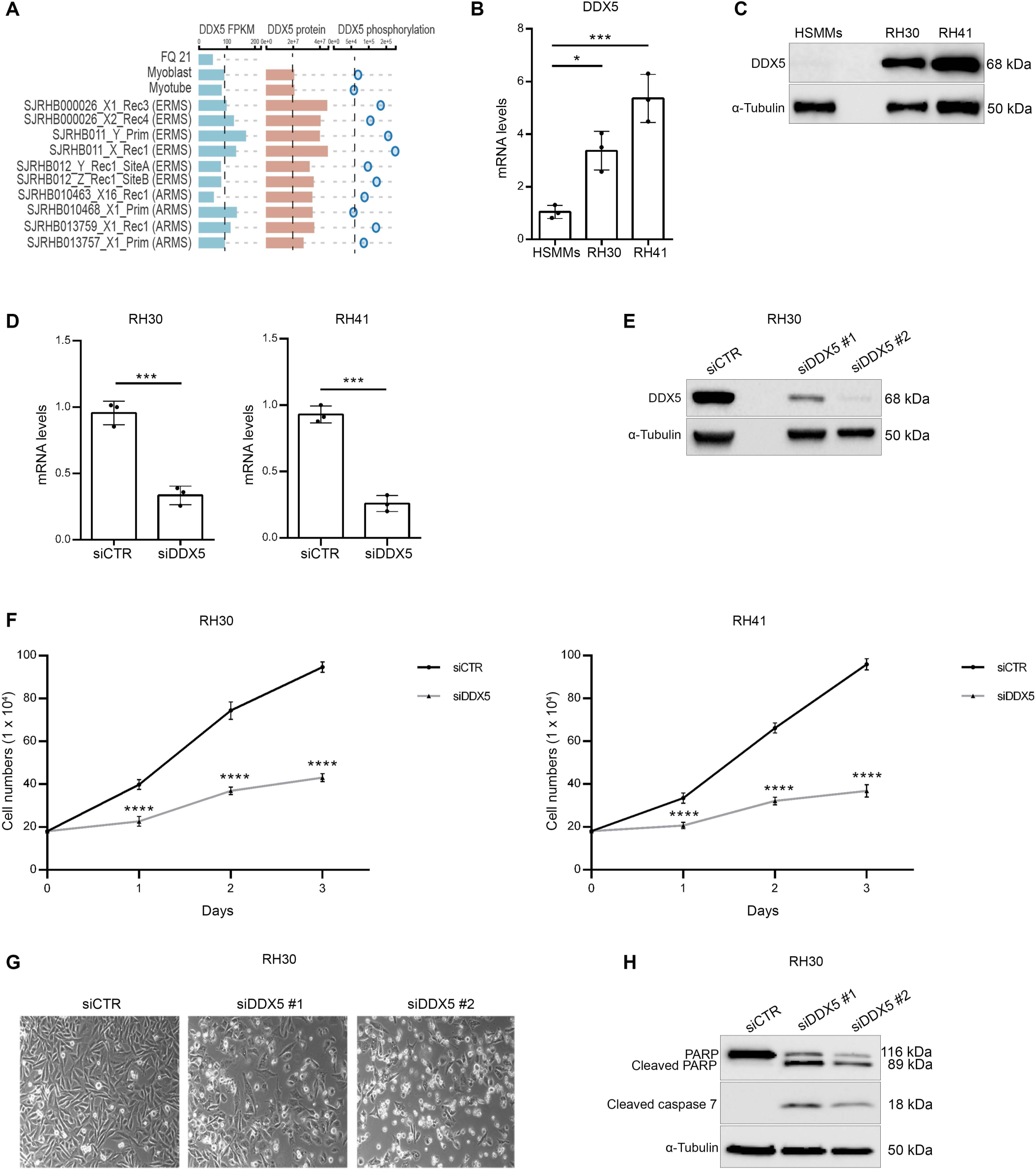
DDX5 is overexpressed in alveolar rhabdomyosarcoma and promotes FP-ARMS survival. **(A)** RNA (FPKM; blue bars) and protein levels (red bars), together with phosphorylation status (blue circles) of DDX5 in orhtotipic RMS patient-derived xenografts (POX), as compared to normal myoblasts and myotubes. Data derive from https://pecan.stjude.cloud/proteinpaint/study/RHB2018 (14). **(B)** Histogram showing the relative mRNA expression levels of DDX5 in RH30 and RH41 cell lines and human skeletal muscle myoblasts (HSMMs). Transcription values were assessed by qRT-PCR and normalized to GAPDH. Graph represents mean+/- SD from n= 3 independent experiments. **(C)** Representative western blot analysis for DDX5 in RH30 and RH41 cell lines and HSMMs. α-tubulin was used as a loading control. **(D) q**RT-PCR analysis of DDX5 mRNA levels in RH30 (left) and RH41 (right) cell lines, after siCTR and siDDX5 treatment. Graphs show mean +/- SD from n= 3 independent experiments. **(E)** Western blot analysis for DDX5 in RH30 cells treated with siCTR, and two different sequences for DDX5 (siDDX5 #1 and #2). Normalization with a-tubulin. **(F)** Cells growth curves after siDDX5 treatment. Cells were counted 1, 2 and 3 days after treatment. Graphs show mean +/- SD from n= 3 independent experiments. **(G)** Representative phase contrast images of RH30 cells 3 days after siDDX5 treatment. **(H)** Western blot analysis for indicated proteins performed on RH30 after siDDX5 and siCTR treatment. α-tubulin was used as control of loading normalization. Statistical significance has been assessed in **(B)** by one-way ANOVA with Bonferroni multiple comparisons test. * p < 0.05; ***p < 0.001; **(D)** by an unpaired Student’s !-test; *** p < 0.001; and **(F)** with 2-way anova with Sidak’s multiple comparison test,. ***p < 0.001 test.

To this end, we inhibited DDX5 expression in FP-ARMS cells through small interfering RNA (siRNA)-mediated knock-down (KD). RH30 and RH41 cells were transfected with two different siRNA against DDX5 or with scramble siRNAs as control (siCTR). As demonstrated by qRT-PCR (**Figure 1D**) and western blot (**Figure 1E**) analysis, both siRNAs efficiently depleted DDX5 in both cell lines. A growth curve performed on both RH30 (**Figure 1F**, *right panel*) and RH41 (**Figure 1F**, *left panel*), demonstrated that DDX5 depletion significantly reduced FP-RMS growth, as compared to control cells (siCTR). An effect that was visible by morphological inspection upon 72 hrs after treatment, (**Figure 1G)**. Western blot analysis for the apoptotic markers cleaved PARP and cleaved caspase 7 (**Figure 1H**), clearly demonstrated the induction of programmed cell death in FP-ARMS upon reduction of DDX5 expression. Taken together, these data led us to conclude that overexpression of DDX5 sustains FP-RMS growth and survival.

### DDX5 promotes AKT signaling stabilizing G9a in FP-RMS

To gain further insights into the mechanism behind DDX5 role in sustaining FP-RMS survival, we performed transcriptional profiling by RNA-seq in siDDX5 Rh30 cells, as compared to siCTR-transfected cells (**Figure 2A**). Notably, enrichment analysis of the differentially expressed genes (DEGs; *p<0*.*05, FC>1*.*5*) found “regulation of RAS protein signal transduction” among categories enriched in down-regulated transcripts (**Figure 2B**). This evidence caught our attention, as RAS pathway is one of the most de-regulated in both FN-RMS and FP-RMS (16). Its predominant downstream signaling pathways, such as the RAF–MEK (mitogen-activated protein kinase (MAPK) kinase–ERK (extracellular signal–regulated kinase) MAPK pathway and the phosphatidylinositol 3-kinase (PI3-kinase)–AKT–mammalian target of rapamycin (mTOR) pathway), are key to maintain cell growth and proliferation, which is why their inhibition is being tested to arrest cancer cell survival (17), with positive effects reported also in RMS (18). Of note, KEGG analysis of ‘RAS signaling’ pathway **(Figure 2C**, *left panel*) indicated that among its downstream cascades, the **‘**Akt signaling’ pathway was the one affected by DDX5 depletion. Accordingly, KEGG on the specific ‘PI3K-Akt signaling’ pathway highlighted downregulation of *AKT* **(Figure 2C**, *right panel*). AKT is involved in cell survival and proliferation, through the mTOR pathway (19), and in apoptosis inhibition by blocking the FOXO cascade (20). To validate this finding, we performed qRT-PCR for AKT on RH30 cell lines after 3 days siDDX5 treatment, demonstrating a significant reduction of AKT mRNA levels as compared to siCTR cells (**Figure 2D**). Moreover, by western blot analysis we demonstrated that *DDX5* silencing induced a significant reduction of the protein levels of AKT and, consequently, of p-AKT, the fully activated form of the kinase phosphorylated on Thr 308 and Ser 473 (**Figure 2E**). These data agree with recent work demonstrating that DDX5 promotes hepatocellular carcinoma cell growth *via* AKT pathway (21), supporting a similar role for DDX5 in sustaining FP-ARMS growth and survival.

**Figure 2.**
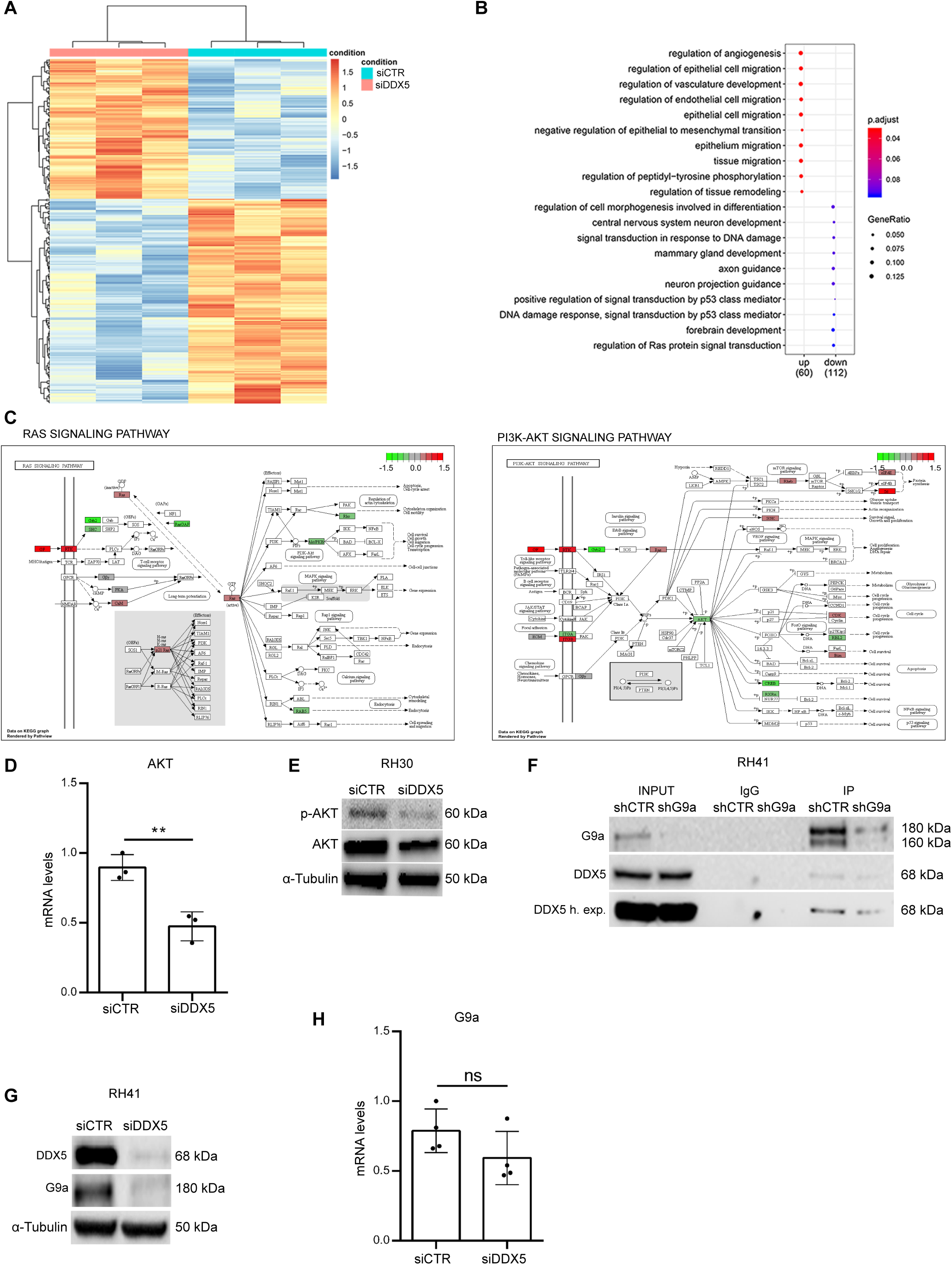
DDX5 interacts with G9a and regulates G9a-AKT signalling. (**A**) Heatmap plot of hierarchical clustering of all differentially expressed genes (DEGs). The X axis represents the two compared samples (siCTR and siDDX5). The Y axis represents DEGs. The color (from blue to red) represents gene expression intensity from low to high. Red indicates upregulated and blue represents downregulated genes. DEGs were selected according to FC > 1,5 e padj < 0.05. (**B**) Dot plot showing the up and down regulated GO terms of biological processes identified. The size of the dot is based on gene count enriched in the pathway, and the color of the dot shows the pathway enrichment significance. (**C**) KEGG analysis of Ras (left) and PI3K-AKT (right) signaling pathways.(**D**) AKT mRNA levels quantified by qRT-PCR after DDX5 silencing in RH30 cells, as compared to siCTR cells. Transcription values were assessed by qRT-PCR and normalized to GAPDH. Graph represents mean +/- SD from n=3 independent experiments. Statistical significance assessed by unpaired Student’s t-test;** p <0.01). (**E**) Western blot analysis of p-AKT and AKT in siCTR and siDDX5 RH30 cells. Normalization with α-tubulin (**F**) Representative western blot analysis of the indicated proteins in G9a immunoprecipitation in nuclear extracts of control (shCTR) and shG9a-treated RH41 cells (last two right lanes). Equal amounts of nuclear extracts were immunoprecipitated with IgG as negative control. Inputs are shown on the left. (**G**) Western blot analysis for G9a and DDX5 in RH30 cells treated with siCTR and siDDX5. α-tubulin was used as loading control. (**H**) qRT-PCR for G9a in siCTR and siDDX5 RH30 cells. Graph shows the mean +/- SD value derived from n=4 independent experiments. Statistical significance has been assessed by an unpaired Student’s t-test; p > 0,05 (no statistical significance, ns).

It has been recently reported that AKT signaling is also pathogenically activated in FP-ARMS by the lysine methyltransferase G9a (22), an evidence that led us to hypothesize that DDX5 and G9a might exert their function *via* a common regulatory axis on AKT. To investigate for this potential cooperation, we performed co-immunoprecipitation studies in RH41 cells in control condition (shCTR) and in cells depleted of G9a (by short-hairpin RNA against G9a, shG9a). As shown in **Figure 2F**, we demonstrated the presence of DDX5 in the immunoprecipitation of G9a, which was reduced in cells with decreased levels of G9a, confirming the specificity of the interaction (**Figure 2F**). In further support for a possible cooperation between G9a and DDX5, we showed that G9a depletion in FP-ARMS cells phenocopied the downregulation of DDX5. Indeed, shG9a FP-ARMS displayed growth arrest (**Supplemental Figure. 2A-B)**; and an increased expression of apoptotic markers (**Supplemental Figure. 2C**). Then, to better investigate the relationship between DDX5 and G9a, we studied their reciprocal modulation. While G9a reduction had no effect on DDX5 protein levels (**Figure 2F**, *input lanes*), we observed that depletion of DDX5 led to a significant reduction of G9a protein in RH41 cells (**Figure 2G**). Since G9a mRNA levels did not significantly decreased after DDX5 silencing (**Figure 2H**), our results suggest that DDX5 regulates G9a post-transcriptionally. In agreement with this idea, it has been recently demonstrated that DDX5 is involved in the alternative splicing of *G9a* transcripts in spermatogonia (23) a mechanism that has been also shown to affect the stability of different G9a isoforms during neuronal differentiation (24). These results lead to speculate that DDX5 might be involved in the control of *G9a* splicing also in FP-ARMS, likely stabilizing specific isoforms and ultimately affecting G9a protein stability.

Taken together, our data indicate that DDX5 function upstream of G9a to promote cell growth and survival, at least in part, via AKT modulation.

### G9a regulates PAX3-FOXO1 protein stability thus sustaining FP-RMS myoblastic state

FP-RMS are addicted to the oncogenic capacity of PAX3-FOXO1, which has been firmly implicated in perpetuating the myoblastic proliferative state of RMS cells (7). In light of our results showing that DDX5 and G9a downregulation disrupt this survival-promoting loop, we decided to investigate whether DDX5 and G9a could be involved in the PAX3-FOXO1 modulation.

Of note, depletion of both *DDX5* (**Figure 3A**) and G9a (**Figure 3B**) caused a marked reduction of PAX3-FOXO1 protein; while PAX3-FOXO1 mRNA levels remained unaffected **(Supplemental Figure. 3A)**, pointing towards a post-transcriptional regulation. Since PAX3-FOXO1 reduction was observed also upon G9a depletion, a condition in which DDX5 levels are unaffected (**Figure 2F**), we hypothesized that G9a is the major regulator of PAX3-FOXO1 oncoprotein in this axis. In agreement with this idea, inhibition of G9a enzymatic activity by treatment with two specific small molecule inhibitors, A366 (**Figure 3C**) and UNC0642 (**Figure 3D**), exerted similar effects on PAX3-FOXO1 levels, further suggesting that G9a mediates PAX3-FOXO1 stability via its enzymatic activity. Accordingly, we found that G9a interacted with PAX3-FOXO1 in two different FP-RMS cell lines (**Figure 3E**), and this interaction was reduced in cells treated with the G9a inhibitor A-366 (**Figure 3F**). As G9a has a well-known role in methylating non-histone proteins, and lysine methylation promotes protein stability (25), these results strongly suggest that G9a might stabilize PAX3-FOXO1 through methylation. In further support of this, transcriptomic analysis by RNA-seq revealed that G9a depletion in FP-ARMS cells (**Supplemental Figure. 3B**) induced transcriptional changes inversely correlated by those imposed by PAX3-FOXO1 expression (26, 27), as revealed by Gene Set Enrichment Analysis (GSEA) (**Figure 3G**). Accordingly, transcript levels of known PAX3-FOXO1 target genes were reduced in G9a KD cells (**Figure 3H**). Further, it has been previously shown that PAX3-FOXO1 activates the RMS master transcription factors (MTFs) MYOD1 and Myogenin, and together with them establishes the epigenome and transcriptional signatures of FP-RMS (7). Strikingly, G9a down-regulation (**Figure 3I**,**L**) and pharmacological inhibition (**Figure 3M**) was sufficient to induce a strong reduction of MYOD1 and Myogenin expression, both at the RNA (**Figure 3I**) and protein (**Figure 3L,M**) levels. This was confirmed by GSEA that revealed a striking negative correlation of transcripts belonging to the “Myogenesis” hallmark in G9a depleted cells (**Supplemental Figure. 3C**), as compared to control. Taken together, these evidences strongly indicate that G9a activity promotes the PAX3-FOXO1-induced myoblastic RMS stage stabilizing the stability of the oncoprotein.

**Figure 3.**
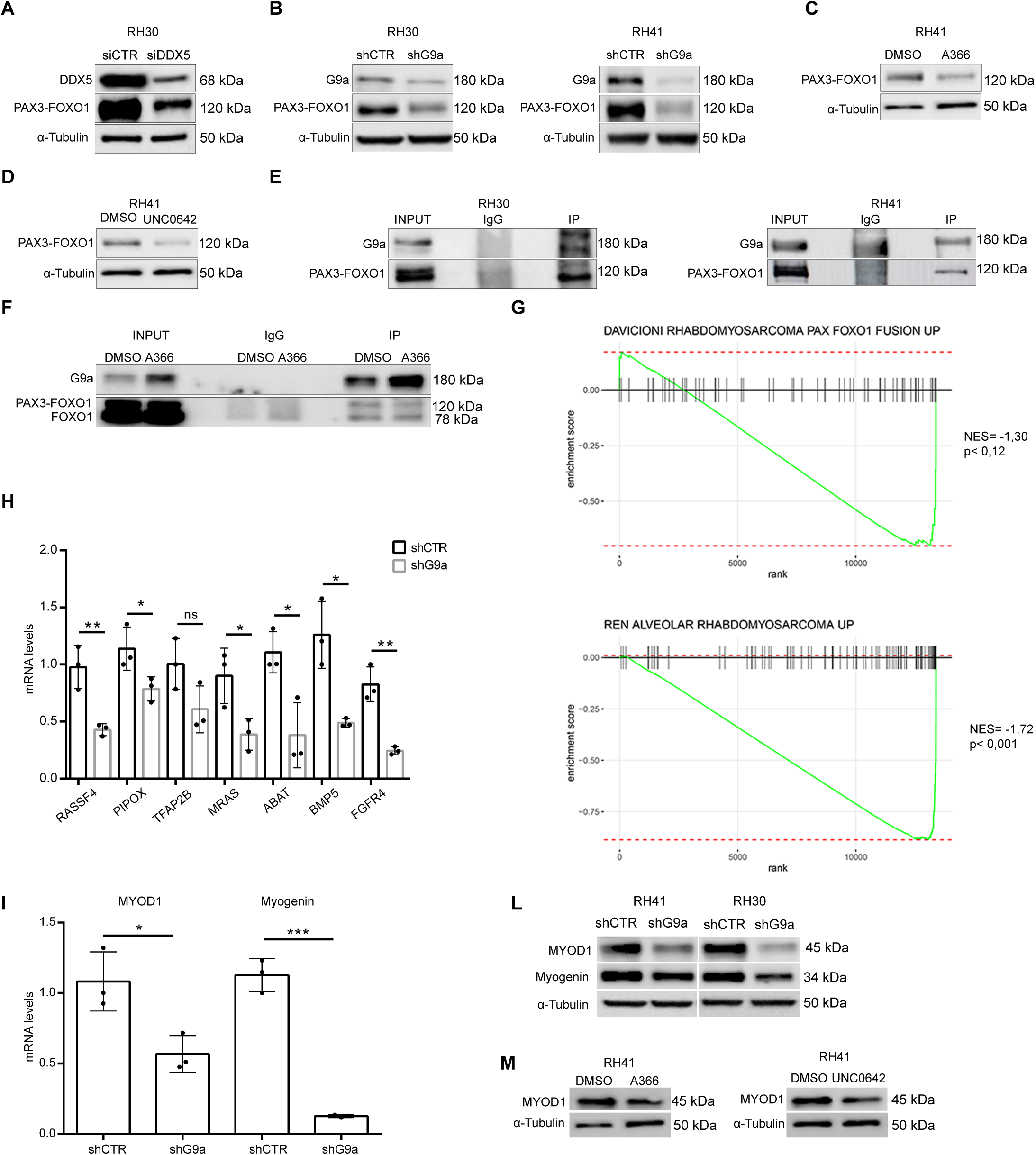
DDX5 and G9a regulate PAX3-FOXO1 expression. (**A**) Western blot analysis for DDX5 and PAX3-FOXO1 in RH30 cells treated with siCTR and siDDX5. α-Tubulin shown as loading control. (**B**) Western blot for G9a and PAX3-FOXO1 in RH30 (left) and RH41 (right) cells treated with shCTR and shG9a. α-Tubulin shown as loading control. (**C-D**) Western blot for PAX3-FOXO1 in RH41 cells treated with 10µM A366 (**C**) and 2 µM UNC0642 (**D**) for six days. Control cells were treated with DMSO and all samples were normalized with α-Tubul-in. (**E**) Western blot analysis of G9a immunoprecipitation in RH30 (left) and RH41 (right) cells. PAX3-FOXO1 is detected in the G9a immunoprecipitates, but not in the IgG negative control. Input are shown on the left. (**F**) Western blot analysis for G9a and PAX3-FOXO1 in RH41 cells treated with A-366 as indicated in (**C**) and immunoprecipitated for G9a. IgG were used as a negative control. (**G**) GSEA of RNA-seq performed in shCTR and shG9a RH41 cells, on genes found upregulated by PAX3-FOXO1, using two different data sets (27) and (26). (**H**) Validation of the indicated PAX3-FOXO1 target genes by qRT-PCR in shCTR and shG9a cells. Transcription values were normalized to GAPDH. Graphs show the mean +/- SD from n=3 independent experiments. (**I**) mRNA levels of MYOD1 and myogenin quantified by qRT-PCR after shG9a treatment in RH41 and RH30 cells. Transcription values were normalized to GAPDH. Graphs show the mean +/- SD from n=3 independent experiments. (**L**) Western blot analysis for MYOD1 and Myogenin in RH41 and RH30 cells after shCTR and shG9a. α-tubulin was used for normalization. (**M**) Western blot of MYOD and MYOG protein levels in RH41 cells treated with 10 µM A366 or with 2 µM UNC0642 for six days. α-tubulin was used for normalization.Statistical significance has been assessed in (**H**) and (**I**) by unpaired Student’s t-test; (* p < 0.05, ** p < 0.01, *** p < 0.001, ns > 0.05).

### DDX5 promotes FP-RMS growth *in vivo*

Our results identified the existence of a three-component regulation axis in which DDX5 functions upstream of G9a and PAX3-FOXO1 to sustain FP-ARMS survival. To unequivocally demonstrate a role of DDX5 in mediating FP-ARMS tumorigenesis *in vivo*, we performed xenografts experiments. To this end, we subcutaneously injected both control (shCtr) and DDX5 depleted RH30 cells (shDDX5) into the flanks of BALB/c Nude mice. Consistent with our *in vitro* data, tumors derived from shDDX5 displayed a significantly reduced growth over time as compared to those derived from control cells (shCtr) (**Figure 4A**); and excised tumors were much smaller than controls (**Figure 4B**). Moreover, immunohistological analysis demonstrated that shDDX5 isolated tumors displayed a significant reduced number of proliferating (Ki67+) cells, as compared to those derived from control (shCtr) cells (**Figure 4C**). These results are consistent with the growth inhibition we previously observed *in vitro* and demonstrate that DDX5 plays a crucial role in FP-RMS growth *in vivo*. Moreover, western blot analysis on xenografts-derived protein extracts confirmed our *in vitro* data, showing a DDX5-dependent expression of G9a and PAX3-FOXO1 in RMS tumors (**Figure 4D-E**), confirming that DDX5 downregulation also reduces pAKT, AKT and MYOD1 protein (**Figure 4E**) levels *in vivo*.

**Figure 4.**
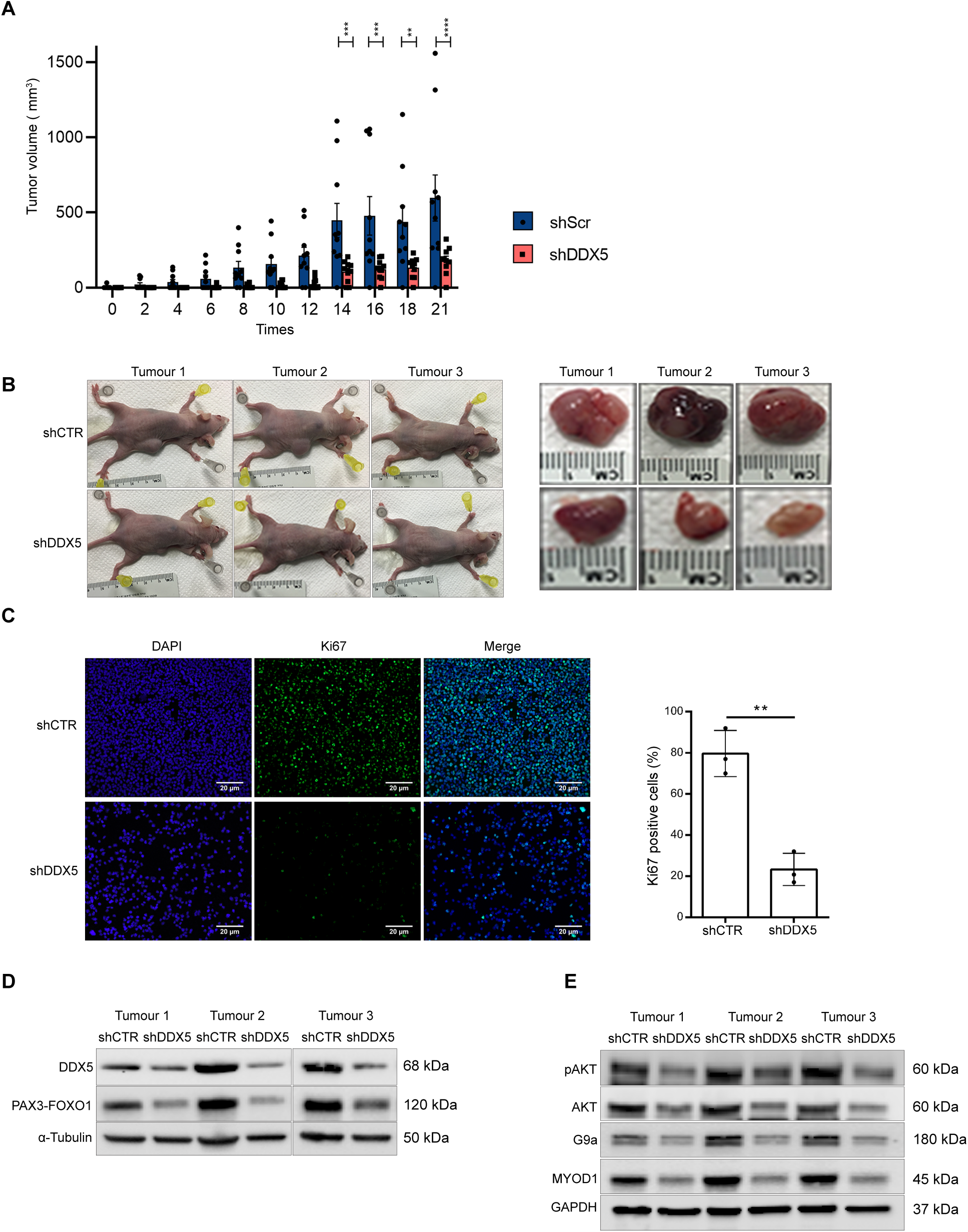
DDX5 promotes FP-RMS growth in vivo. **(A)** Bar charts represent tumor volumes in sCTR and shDDX5 treated mice from day of injection (day 0) to day of tumor resection (day 21). Data are presented as mean +/-sem of n=10 mice/experimental group. Statistical significance assessed by 2-way Anova, with Sidak’s multiple comparison test. ** p<0.01, *** p<0.001, **** p<0.0001. (**B**) Three representative shCTR and shDDX5 treated mice and matching resected tumors at day 21. (**C**) Representative images of Ki67 immunostaining (green) on isolated shCTR and shDDX5 tumors. DAPI (blue) was used to stain nuclei. Scale bars: 20 μm. Histogram (on the right) shows quantification of the percentage of Ki67+ cells. Data are represented as mean +/- sd of n=3 independent tumors. (**D-E**) Western blot analysis for the indicated protein on extracts derived from n=3 tumors/experimental group. α-tubulin and GAPDH were used for normalization.

In sum, our data identify a major role for DDX5 in sustaining FP-ARMS survival and designate it as a possible novel therapeutic target for rhabdomyosarcoma. DDX5 ha

## Methods

### Cell lines

All cell lines were maintained in a humidified incubator at 37°C with 5% CO2. Primary <u>h</u>uman <u>s</u>keletal <u>m</u>uscle <u>m</u>yoblasts (HSMMs) were cultured in growth medium (SkGM-2 Bullet Kit, Lonza). HEK293T cells (kindly gifted by Slimane Ait-Si-Ali lab) for the production of lentiviral particles were cultured in Dulbecco’s modified Eagle’s medium (DMEM) (Sigma-Aldrich, D5671), supplemented with 10% FBS (Corning, 35-015-CV), 2 mM L-glutamine and 100 U/ml penicillin/ streptomycin. ARMS cell lines RH30 and RH41 were kindly provided by Rossella Rota (Bambino Gesù Children’s Hospital, Rome, Italy). RH30 and RH41 were maintained in RPMI 1640 with L-glutamine (Sigma-Aldrich, R8758,) supplemented with with 1% penicillin/streptomycin and 10% FBS (Corning, 35-015-CV). Cells were treated with 2µM UNC0642 and 10 µM A366 (Sigma-Aldrich, SML1410-25MG) Control cells were treated with equivalent concentrations of DMSO (Sigma-Aldrich). Several first passage aliquots of each cell line were stored in liquid nitrogen for subsequent assays.

### Cells transfection

RH30 and RH41 cells were transfected with 100nM of human DDX5 specific siRNA (siDDX5 #1 and siDDX5 #2, Sigma-Aldrich) or scrambled control siRNA (siCTR) (siRNA universal negative control, Sigma-Aldrich) using Lipofectamine 2000 (Invitrogen) according to the manufacturer’s protocol. Transfection with siRNAs was executed when cultured cells reached a confluency of 60% in 6 well plates. Transfection was carried out according to our adapted protocol in RPMI growth medium for 4-6 hours at 37°C. Transfection was then stopped by removing the growth medium and replacing it with RPMI with 10% fetal bovine serum. RNA or protein were isolated 72 h post-transfection for all assays. The targeted sense and antisense strands are shown below: siDDX5 #1 Sense: 5’-AACCGCAACCAUUGACGCCAU-3’ Antisense: 5’-AUGGCGUCAAUGGUUGCGGUU-3’ siDDX5 #2 Sense: 5’-GGCUAGAUGUGGAAGAUGU-3’ Antisense: 5’-ACAUCUUCCACAUCUAGCC-3’

### Short hairpin (sh)RNA lentivirus production and cell infections

Lentiviruses were produced in HEK293T packaging cells seeded in 100 mm culture dishes and transfected in 10ml of DMEM medium, using lipofectamine 2000 (Thermo Fisher Scientific), with lentiviral packaging vectors psPAX2 (7 μg; Addgene) and pMD2.G (3,5 μg; Addgene) and 10 *µ*g lentiviral expression constructs shRNA pLKO.1-puro (G9a Mission shRNA, Sigma-Aldrich, TRCN0000115671, NM_025256,); For DDX5 knockdown the custom sequence AACCGCAACCAUUGACGCCAU (Sigma-Aldrich DDX5 Mission shRNA plasmid DNA, NM_004396.) was cloned in the pLKO.1-puro vector. The non-silencing shCTR (mission control shRNA plasmid DNA) was purchased from Sigma aldrich. Transfection medium was replaced 24 h later with new complete DMEM and 48 h after transfection the lentiviral containing medium was collected, spun to remove cell debris, and the supernatant filtered through a 0.45 μm low protein binding filters. Viral aliquots immediately stored at -80°C. RH30 and RH41 target cells were plated in a 100 mm dish (1,5×10^6^ cells) and, 24 h later, were infected with lentiviral pLKO.1-puro vectors expressing specific shRNA sequences for 24 h in the presence of polybrene (8μg/ml; Sigma-Aldrich). After further 24 hrs, RH30 and RH41 cells were selected with 1 μg/ml puromycin (Sigma-Aldrich, P8833) for 3 days. Cells were harvested at different time points for subsequent experiments. All shRNAs were obtained from Sigma Aldrich.

### In vitro proliferation assays

Cells transfected with siRNA DDX5 and cells transduced with lentivirus shRNA G9a were seeded in 6-well plates (1.8 10^5^ cells per well) and cell proliferation was evaluated by counting trypsinized cultures at 1, 2 and 3 days in RH30 and RH41 cells.

### RNA extraction and Quantitative real time PCR (qRT-PCR)

Cells were harvested and centrifuged at 3000 rpm for 5 min at 4°C. Supernatant was then removed and cell pellet was resuspended in 1 ml of ice-cold PBS and centrifuged at 2000 rpm for 5 min at 4°C. After removing the supernatant, cell pellet was resuspended in 1 mL of TRI Reagent (Sigma aldrich) and RNA extraction was carried out following manufacturer’s protocol. Quantity of RNA samples were assessed with NanoDrop analysis (NanoDrop Technologies). cDNA synthesis was performed using a High Capacity cDNA Reverse Transcription Kits (Applied Biosystems). qRT-PCR was performed with a StepOne plus Real-Time PCR System (Applied Biosystems) to analyze relative gene expression levels using SYBR Green Master mix (Applied Biosystems) following manufacturer indications.

PCR amplification was performed as follows: 95°C 5 minutes, followed by 95°C for 10s, annealing at 60°C for 10s, followed by 45 cycles at 72°C for 10s. Melting curves were generated and tested for a single product after amplification. Expression of each target was calculated using the 2-ΔΔCt method and expressed as a relative mRNA expression. Relative expression values were normalized to the housekeeping gene GAPDH. qRT-PCR was done using reaction duplicates and three independent biological replicates were done for each analysis. Error bars indicate the mean ± standard deviation.

The primers we used are:

DDX5, For: 5’-GCCGGGACCGAGGGTTTGGT-3’, Rev: 5’-CTTGTGCTGTGCGCCTAGCCA-3’; AKT, For: 5’-TCTATGGCGCTGAGATTGTG-3’, Rev: 5’-CTTAATGTGCCCGTCCTTGT-3’; G9a For: 5’-AGAGTGTGGACGGAGAGCTC-3’ Rev: 5’-GGTCTCCCGCTTGAGGAT-3’; MyoD For: 5’-CCGCCTGAGCAAAGTAAATGA-3’; Rev 5’-GCAACCGCTGGTTTGGATT-3’; RASSF4, For: 5’-AGTCCATTCAGAAGTCGGAGC-3’; Rev: 5’-CCCCAGGCAATGTTGAGGAG-3’; PIPOX For: 5’-GGAGCAGTTCTTTCTACCACAC-3’; Rev: 5’ TTCCCAGCAGCAGTAATCCA-3’; TFAP2B, For: 5’-TCAATGCATCTCTCCTCGGC-3’; Rev:5’-CAGCTTCTCCTTCCACCAGG-3’;MRAS,For:TGGCGACCAAACACAATATTCC; Rev: TCTCCCCGCCATTTGGTTTT-3’;ABAT,For:5’-CTGCCTCCGGAGAACTTTGT-3’;Rev:5’-TTTCCTTGCTCCGGTACCAC-3’; BMP5 For: AATGCCACCAACCACGCTAT Rev: 5’-GCCACATGAGCGTACTACCA-3’; FGFR4, For: 5’-TGGCCGTCAAGATGCTCAAA-3’; Rev:5’-GTACAGGGGCCCTTCCTGG-3’;GAPDHFor:5’-TCTGGTAAAGTGGATATTGTTGCC-3’; Rev: 5’-CAAGCTTCCCGTTCTCAGCC-3’;PAX3-FOXO1, For: 5’-AGACAGCTTTGTGCCTCCAT-3’; Rev: 5’-CTCTTGCCTCCCTCTGGATT-3’; myogenin, For: 5’-TCAACCAGGAGGAGCGTGA-3ì; Rev: 5’-TGTAGGGTCAGCCGTGAGCA-3’

### Protein extraction and Western blotting

Cells were harvested and centrifuged at 3000 rpm for 5 min at 4°C. Supernatant was then removed and cell pellet was resuspended in 1 ml of ice-cold PBS and centrifuged at 2000 rpm for 5 min at 4°C. After removing the supernatant, cell pellet was resuspended in lysis RIPA buffer supplemented with protease inhibitor cocktail and phosphatase inhibitors (Roche) and incubated in ice for 30 min. Samples were then sonicated in a water bath for 10 min (30 sec ON/ 30 sec OFF) and centrifuged at 15000 rpm for 15 min at 4°C. Supernatant was then transferred in a new tube and proteins were quantified by BCA assay (Thermo Fisher Scientific) according to the manufacturer’s protocol. Cell lysates were resolved on 4%-20% Mini-PROTEAN TGX gels (Bio-Rad Laboratories) and then transferred to nitrocellulose membrane (Amersham) using Trans-Blot Turbo Transfer system (Bio-Rad Laboratories). Membranes were blocked with 5% nonfat dried milk in Tris-buffered saline/Tween (TBS-T; 0.1%) for 1 hours at room temperature with gentle shaking, followed by overnight incubation at 4°C with various antibodies. The primary antibodies we used were: DDX5 (Cell Signalling, #9877), G9a (Cell signalling, #3306), FOXO1 (Cell Signalling, #2880), p-AKT (Ser473) (Cell signaling, #4058), AKT (Cell signaling #4685) PARP (Cell signaling #9542), cleaved caspase 7 (Cell signaling #8438), cleaved caspase 9 (Cell signaling #9505), MYOD (Santa Cruz Biotechnology, C-20), myogenin (DSHB, F5D), GAPDH (Sigma, G9545), α-tubulin (Sigma-Aldrich, T5168). Membranes were then incubated with HRP-conjugated secondary antibodies (IgG-HRP Santa Cruz Biotechnologies) for 1 hour at RT and after incubation the blots were developed in an ECL detection solution (Clarity Max ECL substrate, Bio-Rad Laboratories) and signal was detected using ChemiDoc (BioRad Laboratories).

### Mouse xenograft experiments

All animal procedures were approved by Italian Ministry of Health and Istituto Superiore di Sanità; approval number 7FF2C.7-EXT.9.

Female Balb/c nude mice (6/7 weeks old) were obtained from (Envigo) and maintained under specific pathogen–free conditions in a temperature- and humidity-controlled environment (Allevamenti Plaisant). 2 × 10^6^ shCTR and shDDX5 RH30 cells were injected subcutaneously into the flank of and, once tumors were palpable, they were measured every other day by measuring 2 diameters (d1 and d2) in right angles using a digital caliper. Total tumor volumes were then calculated by the formula V = (4/3)πr3; r = (d1 + d2)/4. On day 21 after the injections, mice were euthanized and resected tumors were fixed in formalin or immersed in liquid nitrogen and stored at -80 degrees. For western analysis tumors were disrupted with a mortar and pestle, followed by sonication in RIPA buffer supplemented with proteinase and phosphatase inhibitors (Roche). Formalin-fixed tumor tissues were embedded in paraffin and and sections were stained with hematoxylin and eosin using standard techniques (data not shown). Tissue sections were deparaffinized, rehydrated, and heated at 95°C for 20 min in pH 6 antigen retrieval buffer. Slides were blocked and incubated with Ki67 antibody (Abcam 15580) overnight at 4°C, then incubated with the secondary antibody (Alexa Fluor 488, Thermo Fisher Scientific). Nuclei were counterstained with DAPI (Sigma). Images were acquired using Axio Observer 443 microscope (ZEISS) and analyzed by ZEN 3.0 (Blue edition) software.

### Co-immunoprecipitation

Co-immunoprecipitation was carried out through magnetic separation. RH30 and RH41 cells were centrifuged at 1200 rpm for 5 min at 4°C, resuspended in lysis buffer (10 mM Tris pH 8, 10 mM NaCl, 0.1 mM EDTA pH 8, 0.1 mM EGTA) with protease inhibitor (Roche) and then incubated on ice for 30 min. A dounce homogenizer was used to mechanically help cell lysis. 10% NP-40 was added to a final concentration of 0,5% and then samples were vortexed and incubated on ice 2-3 min. Samples were then centrifuged at 4000 rpm for 5 min at 4°C in order to pellet nuclei. Nuclei were resuspended in nuclei lysis buffer (20 mM Tris pH 8, 400 mM NaCl, 1 mM EDTA pH 8, 1 mM EGTA) with protease inhibitor (Roche) and incubated for 10 min on ice to increase lysis efficiency. Lysates were then sonicated for 10 min (30 sec On/30 sec OFF at high intensity) and then centrifuged at top speed for 20 min at 4°C. Supernatant, containing nuclear extract (NE), was then transferred into a new tube: 50 µl on NE was saved to use as input and to quantify NE concentration. NE was then precleared with 10 µl protein A/G magnetic beads (Thermo Fisher Scientific), washed with IP lysis buffer (50 mM Tris pH 8, 150 mM NaCl, 1 mM EDTA pH 8, 1 mM EGTA) for 2 hrs at 4°C on rotating wheel.

After preclearing, NE were diluted 1:5 in IP buffer with protease inhibitor (Roche). NE was incubated overnight at 4°C on rotating wheel with 10 ug of G9a antibody (Abcam, ab185050). The following day we added pre-blocked protein A/G magnetic beads (Thermo Fisher Scientific) to each sample and incubated for 2 hrs at 4°C on rotating wheel. Samples were then washed six times, one wash every 5 min, with IP buffer. At the end, beads were separated on a magnet and the immunocomplexes were resuspended with IP buffer and LSB buffer (Biorad Laboratories) for further analysis.

### RNA-Sequencing

#### RNAseq in sIDDX5 FP-RMS

Total RNA was extracted and quantified as previously described. RNA-Seq libraries preparation and sequencing was performed by the IGA Technology Services (Udine, Italy) using the Illumina TruSeq Stranded mRNA Kit (Illumina, San Diego, CA) according to manufacturer’s instructions. The final libraries for paired-end sequencing of 150 base pairs were carried out on an Illumina NovaSeq6000 (Illumina, San Diego, CA) with an average of 55 million of reads per sample. Processing raw data for both format conversion and de-multiplexing were performed by *Bcl2Fastq* version 2.20 of the Illumina pipeline. Reads quality was evaluated using *FastQC* (version 0.11.8, Babraham Institute Cambridge, UK) tool, then adapter sequences were masked with *Cutadapt* version 1.11 from raw fastq data using the following parameters: *--anywhere* (on both adapter sequences) *--overlap 5 --times 2 --minimum-length 35 --mask-adapter*.

Reads were mapped to the human Ensembl GRCh38 transcriptome index (release 96) using *kallisto* (version 0.46.0) (28). The following flags were used for *kallisto*: *-b 30 --bias*. Gene-level normalization and differential gene expression analysis were performed using Bioconductor (29) R (version 3.6.2) (R Core Team, 2015) package *DESeq2* version 1.26 (30) accounting for the presence of batch effects. The figures were obtained using the *R* environment with package *ggplot2* version 3.3.0 and *pheatmap* version 1.0.12.

### RNAseq in shG9a FP-RMS

RNA-seq libraries from total RNA (100 ng) from each sample were prepared using QuantSeq 3’mRNA-Seq Library prep kit (Lexogen, Vienna, Austria) according to manufacturer’s instructions, at Telethon Institute of Genetics and Medicine (TIGEM). The amplified fragmented cDNA of 300 bp in size were sequenced in single-end mode using the NextSeq500 (Illumina) with a read length of 75 bp. Reads quality was evaluated using *FastQC* (version 0.11.8, Babraham Institute Cambridge, UK) tool and was trimmed using *TrimGalore* software to remove adapter and low-quality bases (Q < 20). Then reads were mapped to the human Ensembl GRCh38 build reference genome using *STAR* version 2.5.0a (31) using Gene annotations corresponding to the Ensembl annotation release 96 which was used to build a transcriptome index and provided to *STAR* during the alignment.

The same gene annotations were used to quantify the gene-level read counts using *HTSeq-count* version 0.8.0 (32) script, subsequently the data normalization and differential analysis for gene expression were performed using Bioconductor (29) R package *edgeR* version 3.28 (33).

### Gene set enrichment analysis

In order to understand biological meaning of the differentially expressed genes the resulting filtered (Benjamini-Hochberg false discovery rate (FDR) adjusted for multiple hypothesis testing p-value < 0.05) genes were clustered by functional annotation using Bioconductor *R* package *clusterProfiler* version 3.14 (34) with annotation of Gene Ontology Database (35) and with annotation of Kyoto Encyclopedia of Genes and Genomes (KEGG) (36) for pathways. Gene Set Enrichment Analysis (GSEA) (37) with pre-ranked, “classic” mode with 10,000 permutations was used to assess the enrichment of the gene profile of siDDX5 or shG9a samples compared to control samples in the curated “hallmark” and “C2” gene set collections (BROAD molecular signature database, MSigDb version 6.2).

## Data availability

RNA-Seq data accompanying this paper are available through NCBI’s Gene Expression Omnibus (GEO) repository, under accession number GSE152358 and GSE152359.

## Statistical analysis

Data were analyzed using Prism (version 6.0; GraphPad Software Inc.), and images were compiled in Photoshop (version 6.0; Adobe Systems). Results are presented as mean ± SD from at least 3 independent experiments. Statistical analysis was conducted using an unpaired Student’s *t*-test, one-way ANOVA or 2-way ANOVA. *P* value of less than 0.05 was considered statistically significant. \**P*< 0.05; \*\**P*< 0.01; \*\*\**P*< 0.001; \*\*\*\**P*< 0.0001.

## Author Contributions

A.G. performed all the experiments, collected and analysed data and prepared figures. V.L. performed bioinformatic analyses. C.M. conceived, supervised the project and wrote the manuscript. All authors discussed results, reviewed and edited the manuscript.

## Acknowledgements

This work was supported by the Italian Association for Cancer Research (AIRC; MyFIRST grant n.18993) and the CNCCS (Collection of National Chemical Compounds and Screening Center). We would like to thank Marco Pezzullo and Cristiano De Stefanis (core facilities, Bambino Gesù Children’s Hospital, Rome, Italy) for the technical FFPE tissues preparation.

---

**Supplemental Figure 1.**
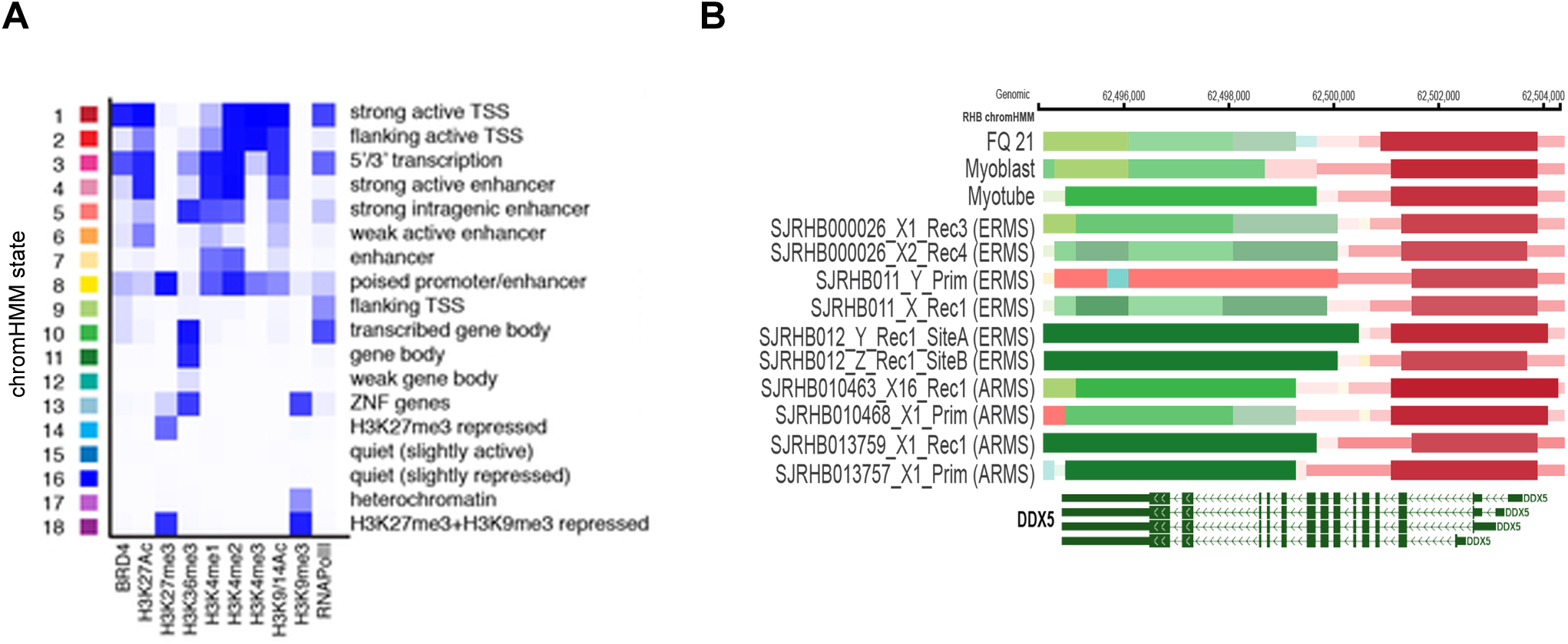
**(A)** Legend of the 18 chromatin hidden Markov modeling (chromHMM) states (15) identified in the study (14). (**B**) ChromHMM state of DDX5 in RMS patient-derived xenografts (PDX) from ERMS and ARMS, as compared to normal myoblasts and myotubes. The bars show a strong active transcription start site (TSS) (red bars) and an actively transcribed gene body (green bars) in either normal myoblasts and myotubes, and primary ERMS and ARMS samples.

**Supplemental Figure 2.**
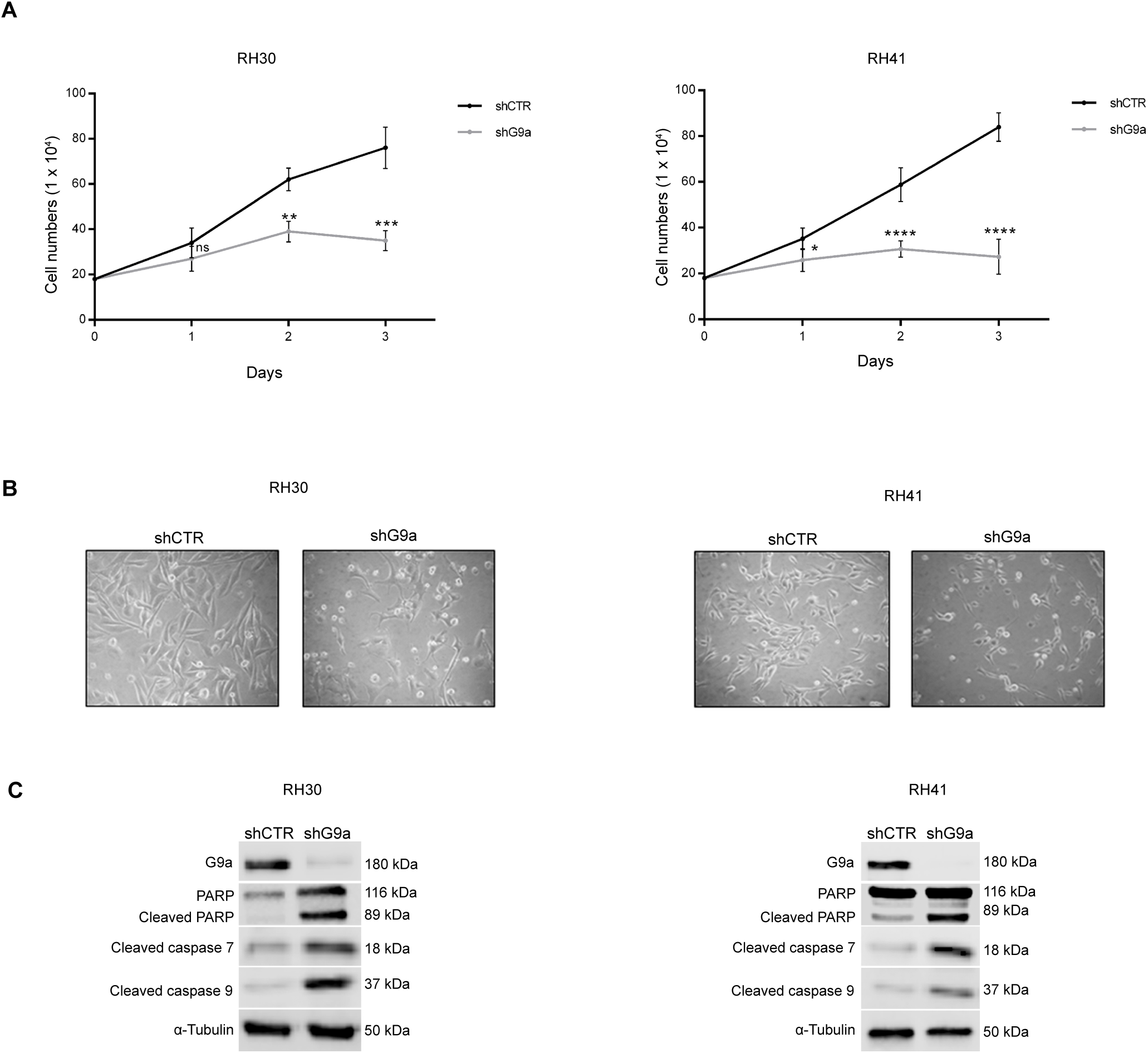
**(A)** RH30 and RH41 cells growth curves after shCTR and shG9a treatment. Cells were counted 1, 2 and 3 days after treatment. Graphs show mean +/- SD from n=3 independent experiments Statistical significance has been assessed by 2-way Anova, with Sidak’s multiple comparison test. ** p<0.01, *** p<0.001, **** p<0.0001. (**B**) Representative phase contrast images of RH30 (left) and RH41 (right), shCTR and shG9a, analyzed after 3 days after treatment. (**C**) Western blot of G9a, cleaved PARP, cleaved caspase 7 and cleaved caspase 9 from extracts of RH30 and RH41 cells 3 days after shG9a silencing. Normalization with α-tubulin.

**Supplemental Figure 3.**
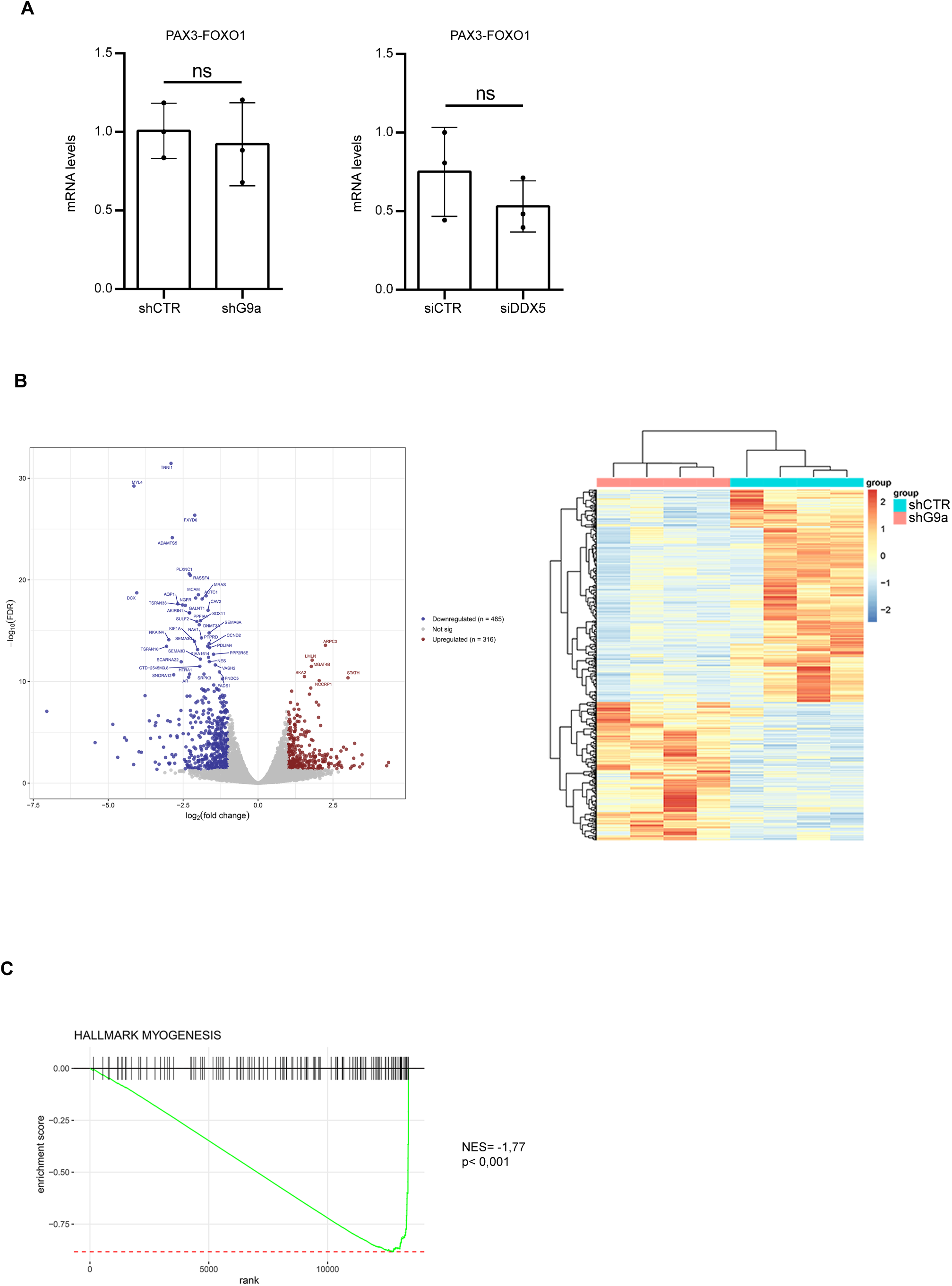
(**A**) qRT-PCR of PAX3-FOXO1 in shG9a and siDDX5 treatments. Graphs show the mean +/-SD values from n=3 independent experiments. Statistical significance has been assessed by an unpaired Student’s t-test; p > 0,05 (no statistical significance, ns). (**B**) RNA-seq analysis after G9a knockdown. Volcano plot of shG9a vs shCTR in RH41 cells (left) showing differentially expressed genes (DEGs) (downregulated genes are highlighted in blue and upregulated genes are red). Heatmap of DEGs in RH41 cells 72h after transfection (right). RNAseq analysis was performed for genes that are regulated by G9a at the transcript-level with more than two-fold regulation (Foldchange > 2 and FDR < 0.05). (**C**) GSEA of RNA-seq performed in shCTR and shG9a RH41 cells, on “Myogenesis” signature.

## References

1. Hettmer S, Li Z, Billin AN, Barr FG, Cornelison DD, Ehrlich AR, et al. Rhabdomyosarcoma: current challenges and their implications for developing therapies. Cold Spring Harb Perspect Med. 2014;4(11):a025650.

2. El Demellawy D, McGowan-Jordan J, de Nanassy J, Chernetsova E, and Nasr A. Update on molecular findings in rhabdomyosarcoma. Pathology. 2017;49(3):238–46.

3. Wachtel M, and Schafer BW. PAX3-FOXO1: Zooming in on an “undruggable” target. Semin Cancer Biol. 2018;50:115–23.

4. Seki M, Nishimura R, Yoshida K, Shimamura T, Shiraishi Y, Sato Y, et al. Integrated genetic and epigenetic analysis defines novel molecular subgroups in rhabdomyosarcoma. Nat Commun. 2015;6:7557.

5. Kohsaka S, Shukla N, Ameur N, Ito T, Ng CK, Wang L, et al. A recurrent neomorphic mutation in MYOD1 defines a clinically aggressive subset of embryonal rhabdomyosarcoma associated with PI3K-AKT pathway mutations. Nat Genet. 2014;46(6):595–600.

6. Shern JF, Chen L, Chmielecki J, Wei JS, Patidar R, Rosenberg M, et al. Comprehensive genomic analysis of rhabdomyosarcoma reveals a landscape of alterations affecting a common genetic axis in fusion-positive and fusion-negative tumors. Cancer Discov. 2014;4(2):216–31.

7. Gryder BE, Yohe ME, Chou HC, Zhang X, Marques J, Wachtel M, et al. PAX3-FOXO1 Establishes Myogenic Super Enhancers and Confers BET Bromodomain Vulnerability. Cancer Discov. 2017;7(8):884–99.

8. Linder P, and Jankowsky E. From unwinding to clamping - the DEAD box RNA helicase family. Nat Rev Mol Cell Biol. 2011;12(8):505–16.

9. Caretti G, Schiltz RL, Dilworth FJ, Di Padova M, Zhao P, Ogryzko V, et al. The RNA helicases p68/p72 and the noncoding RNA SRA are coregulators of MyoD and skeletal muscle differentiation. Dev Cell. 2006;11(4):547–60.

10. Dardenne E, Polay Espinoza M, Fattet L, Germann S, Lambert MP, Neil H, et al. RNA helicases DDX5 and DDX17 dynamically orchestrate transcription, miRNA, and splicing programs in cell differentiation. Cell Rep. 2014;7(6):1900–13.

11. Yang L, Lin C, and Liu ZR. Phosphorylations of DEAD box p68 RNA helicase are associated with cancer development and cell proliferation. Mol Cancer Res. 2005;3(6):355–63.

12. Shin S, Rossow KL, Grande JP, and Janknecht R. Involvement of RNA helicases p68 and p72 in colon cancer. Cancer Res. 2007;67(16):7572–8.

13. Nyamao RM, Wu J, Yu L, Xiao X, and Zhang FM. Roles of DDX5 in the tumorigenesis, proliferation, differentiation, metastasis and pathway regulation of human malignancies. Biochim Biophys Acta Rev Cancer. 2019;1871(1):85–98.

14. Stewart E, McEvoy J, Wang H, Chen X, Honnell V, Ocarz M, et al. Identification of Therapeutic Targets in Rhabdomyosarcoma through Integrated Genomic, Epigenomic, and Proteomic Analyses. Cancer Cell. 2018;34(3):411–26 e19.

15. Ernst J, and Kellis M. ChromHMM: automating chromatin-state discovery and characterization. Nat Methods. 2012;9(3):215–6.

16. Skapek SX, Ferrari A, Gupta AA, Lupo PJ, Butler E, Shipley J, et al. Rhabdomyosarcoma. Nat Rev Dis Primers. 2019;5(1):1.

17. Cox AD, Fesik SW, Kimmelman AC, Luo J, and Der CJ. Drugging the undruggable RAS: Mission possible? Nat Rev Drug Discov. 2014;13(11):828–51.

18. Yohe ME, Gryder BE, Shern JF, Song YK, Chou HC, Sindiri S, et al. MEK inhibition induces MYOG and remodels super-enhancers in RAS-driven rhabdomyosarcoma. Sci Transl Med. 2018;10(448).

19. Laplante M, and Sabatini DM. mTOR signaling in growth control and disease. Cell. 2012;149(2):274–93.

20. Zhang X, Tang N, Hadden TJ, and Rishi AK. Akt, FoxO and regulation of apoptosis. Biochim Biophys Acta. 2011;1813(11):1978–86.

21. Xue Y, Jia X, Li L, Dong X, Ling J, Yuan J, et al. DDX5 promotes hepatocellular carcinoma tumorigenesis via Akt signaling pathway. Biochem Biophys Res Commun. 2018;503(4):2885–91.

22. Bhat AV, Palanichamy Kala M, Rao VK, Pignata L, Lim HJ, Suriyamurthy S, et al. Epigenetic Regulation of the PTEN-AKT-RAC1 Axis by G9a Is Critical for Tumor Growth in Alveolar Rhabdomyosarcoma. Cancer Res. 2019;79(9):2232–43.

23. Legrand JMD, Chan AL, La HM, Rossello FJ, Anko ML, Fuller-Pace FV, et al. DDX5 plays essential transcriptional and post-transcriptional roles in the maintenance and function of spermatogonia. Nat Commun. 2019;10(1):2278.

24. Fiszbein A, Giono LE, Quaglino A, Berardino BG, Sigaut L, von Bilderling C, et al. Alternative Splicing of G9a Regulates Neuronal Differentiation. Cell Rep. 2016;14(12):2797–808.

25. Cornett EM, Ferry L, Defossez PA, and Rothbart SB. Lysine Methylation Regulators Moonlighting outside the Epigenome. Mol Cell. 2019;75(6):1092–101.

26. Ren YX, Finckenstein FG, Abdueva DA, Shahbazian V, Chung B, Weinberg KI, et al. Mouse mesenchymal stem cells expressing PAX-FKHR form alveolar rhabdomyosarcomas by cooperating with secondary mutations. Cancer Res. 2008;68(16):6587–97.

27. Davicioni E, Finckenstein FG, Shahbazian V, Buckley JD, Triche TJ, and Anderson MJ. Identification of a PAX-FKHR gene expression signature that defines molecular classes and determines the prognosis of alveolar rhabdomyosarcomas. Cancer Res. 2006;66(14):6936–46.

28. Bray NL, Pimentel H, Melsted P, and Pachter L. Near-optimal probabilistic RNA-seq quantification. Nat Biotechnol. 2016;34(5):525–7.

29. Gentleman RC, Carey VJ, Bates DM, Bolstad B, Dettling M, Dudoit S, et al. Bioconductor: open software development for computational biology and bioinformatics. Genome Biol. 2004;5(10):R80.

30. Love MI, Huber W, and Anders S. Moderated estimation of fold change and dispersion for RNA-seq data with DESeq2. Genome Biol. 2014;15(12):550.

31. Dobin A, Davis CA, Schlesinger F, Drenkow J, Zaleski C, Jha S, et al. STAR: ultrafast universal RNA-seq aligner. Bioinformatics. 2013;29(1):15–21.

32. Anders S, Pyl PT, and Huber W. HTSeq--a Python framework to work with high-throughput sequencing data. Bioinformatics. 2015;31(2):166–9.

33. Robinson MD, McCarthy DJ, and Smyth GK. edgeR: a Bioconductor package for differential expression analysis of digital gene expression data. Bioinformatics. 2010;26(1):139–40.

34. Yu G, Wang LG, Han Y, and He QY. clusterProfiler: an R package for comparing biological themes among gene clusters. OMICS. 2012;16(5):284–7.

35. Ashburner M, Ball CA, Blake JA, Botstein D, Butler H, Cherry JM, et al. Gene ontology: tool for the unification of biology. The Gene Ontology Consortium. Nat Genet. 2000;25(1):25–9.

36. Ogata H, Goto S, Sato K, Fujibuchi W, Bono H, and Kanehisa M. KEGG: Kyoto Encyclopedia of Genes and Genomes. Nucleic Acids Res. 1999;27(1):29–34.

37. Subramanian A, Tamayo P, Mootha VK, Mukherjee S, Ebert BL, Gillette MA, et al. Gene set enrichment analysis: a knowledge-based approach for interpreting genome-wide expression profiles. Proc Natl Acad Sci U S A. 2005;102(43):15545–50.

